# How often do Protein Genes Navigate Valleys of Low Fitness?

**DOI:** 10.1101/592444

**Authors:** Erik D. Nelson, Nick V. Grishin

## Abstract

In order escape from local fitness peaks, a population must navigate across valleys of low fitness. How these transitions occur, and what role they play in adaptation, have been subjects of active interest in evolutionary genetics for almost a century. However, to our knowledge, this problem has never been addressed directly, by considering the evolution of a gene, or group of genes, as a whole, including the complex effects of fitness interactions among multiple loci. Here, we use a precise model of protein fitness to compute the probability *P*(*s*, Δ*t*) that an allele, randomly sampled from a population at time *t*, has crossed a fitness valley of depth *s* during an interval [*t* − Δ*t, t*] in the immediate past. We study populations of model genes evolving under equilibrium conditions consistent with those in mammalian mitochondria. From this data, we estimate that genes encoding small protein motifs navigate fitness valleys of depth 2*Ns* ≳ 30 with probability *P* ≳ 0.1 on a time scale of human evolution, where *N* is the (mitochondrial) effective population size. The results are consistent with recent findings for Watson–Crick switching in mammalian mitochondrial tRNA molecules.

## 1. Introduction

The effect of a mutation on the fitness of an organism usually depends on the genetic background, or context in which it occurs, a phenomenon known as epistasis [1,2]. Because of this, the fitness landscape of a gene, a group of genes, or an organism will contain many isolated peaks and valleys [3,4], resembling the energy landscape of a physical system such as a glass. Under selection pressure, a population tends to evolve along a path of steepest ascent in fitness until it arrives in the neighborhood of a local fitness peak; In order to escape a sub–optimal fitness peak, the population, or some part of the population, must traverse across a valley of lower fitness. How such transitions occur [1,4–8], and how they relate to adaptation [9] have remained subjects of active interest in evolutionary genetics for almost a century.

The most basic example of valley crossing is realized in the compensatory interaction of individually deleterious mutations at two genetic loci – for instance, as might result from the physical interaction between amino acids in a protein, or a pair of nucleotides in an RNA molecule [5]. The archetypal model of this situation consists of a pair of diallelic loci with initial and final states AB and A′B′ respectively; Mutations to A′ and B′ incur a fitness cost s relative to AB when introduced individually, but are neutral when introduced jointly. Kimura was the first to study this problem using the diffusion approach [5,10], and he found that deep fitness valleys could be crossed on a relatively short time scale if mutation rates are sufficiently large – specifically, when 2*Nμ* = 1 where *N* is the population size and *μ* is the mutation rate per gene per generation. In this case, fitness valleys are navigated by a process known as stochastic tunneling [11], in which a small fraction of genes accumulate in an intermediate state, are compensated by a second mutation, and ultimately proceed to fixation – the intermediate acting as a kind of stepping stone [12]. The situation studied by Kimura closely resembles the process of Watson–Crick switching between favorably paired nucleotides in RNA stem sites, and in particular, switching in mammalian mitochondrial (mt) tRNA molecules where stochastic tunneling is significant. Meer et al. investigated this problem somewhat recently [13], and, using Kimura’s model, they found that mammalian mt tRNA switches may navigate valleys of depth even as large as 2*Ns* ≃ 50 (here, we assume that, for equal numbers of males and females, the effective population size for mitochondrial genes is one fourth the effective population size for nuclear genes [14,15]). In support of this result, Meer et al. obtain essentially the same estimate for 2*Ns* from the frequency (*p*) of disrupted Watson–Crick pairs using the relation *p* = *μ/s* for mutation–selection balance [16]. To put this number into context, it is at least ten times larger than would be expected if the same model had evolved by sequential fixation of deleterious and compensatory mutations (i.e., as would be expected when *μN* ≪ 1 [17]).

While these estimates may be accurate, it is difficult to reconcile the evolutionary dynamics of folded biomolecules with two–locus models. Naturally evolving genes encoding proteins and RNA molecules are always faced with a complex spectrum of possible routes on their fitness landscapes, and it is these spectra that ultimately determine the rate for crossing valleys of a given depth. Even for tRNA molecules, compensation of disrupted Watson–Crick pairs seems to occur more often through complex, indirect mechanisms than through direct compensation to restore Watson–Crick pairing [18]. Proteins are more connected objects than RNA molecules (i.e., with more opportunities for epistatic interactions between loci), and the greater complexity of protein sequences is almost certain to present a more complex spectrum of possible routes to a protein gene in which valleys (ravines, etc.) are entered and exited in multiple steps (Figure A1).

Are the large effects predicted by Meer et al. common in biomolecular evolution? To our knowledge, this kind of question has never been asked directly, by considering the problem of valley crossing for a protein or RNA molecule as a whole, including the complex effects of fitness interactions between multiple loci. Here, we simulate the evolution of a small protein motif using an exact fitness model that is simple enough to allow for adequate sampling of valley crossing statistics. We evolve populations of model genes by haploid Wright–Fisher dynamics across a range of mutation rates spanning the sequential fixation (*μN* ≪ 1) and stochastic tunneling (*μN* ≥ 1) regimes, and we record the mutational paths of all alleles in our populations. Using this data, we compute the probability *P*(*s*, Δ*t*) that an allele, randomly sampled from a population at time *t*, has crossed a fitness valley of depth *s* during a time interval [*t* − Δ*t, t*] in the immediate past. Surprisingly, we find that, even on the time scale of human evolution, genes encoding small protein motifs evolving under conditions consistent with mammalian mitochondria already navigate fitness valleys of depth 2*Ns* ≳ 30 with probability *P* ≳ 0.1, in rough agreement with the estimate for Watson–Crick switching in mt tRNAs provided by Meer et al.

## 2. Methods

Epistatic effects play an essential part in protein evolution [19–24], and because these effects depend on the relative probabilities of conformations in protein ensembles [25–27], it is important to select a model in which the salient properties of protein ensembles are retained as much as possible. Ultimately, we found that we could obtain sufficient data for valley crossing statistics in a reasonable period of time using small lattice proteins. Lattice models have been used extensively in studies of protein folding and evolution, and the model we employ here is similar to one recently used to explore the effects of epistasis on the predictability of protein evolutionary pathways [25].

Below, we evolve lattice proteins under equilibrium conditions to maintain marginal stability in a specific folded (native) conformation. The stability of a protein is measured, as usual, by the free energy difference between the native conformation and the rest of the conformational ensemble,

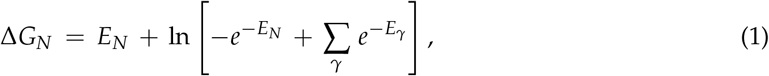

where *E_γ_* is the energy of conformation *γ*, the subscript *N* denotes the native conformation, and factors of temperature are absorbed into the definition of energy; The energy of a conformation is determined from its amino acid contacts by empirical amino acid contact potentials [28] (as a result, energies are defined in units of *RT* ≃ 0.6 kcal / mol).

We assume that mis–folded proteins are non–functional, and otherwise toxic to an organism [29–31]. In this case, protein fitness can be defined by the probability of finding an individual protein folded in its native conformation [25,32],

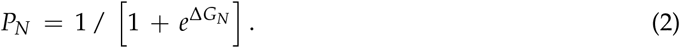

However, since most naturally occurring proteins are only marginally (as opposed to maximally) stable [24], we decided to model fitness using a logistic function

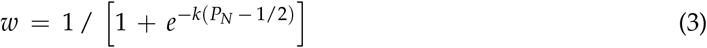

where *k* = 15 (Figure S1). Under this condition, evolved genes in our simulations typically encode proteins with *P_N_* > 0.75 or, equivalently, Δ*G_N_* < −1 [33].

We evolve populations of protein genes by plain Wright–Fisher dynamics, with discrete generations, fixed population size, and no recombination [34]. In each generation, a Poisson random number of nucleotide sites with mean *μN* are selected at random; The sites are subjected to random mutation, the fitness values of mutant alleles are computed, and *N* offspring are selected from the population to form the next generation. The probability that an allele *i* survives to the next generation is 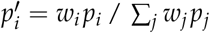, where *p_i_* is the frequency of allele *i* in the current generation [35].

In the absence of recombination, each allele in a population has a unique mutational history extending back to the origin of a simulation. To describe the statistics of valley crossing, we record the histories of all alleles, and we compute *P*(*s*, Δ*t*), the probability that an allele, randomly sampled from the population at time *t*, has crossed a fitness valley of depth *s* during the time interval [*t* − Δ*t, t*], where time is measured in generations. Depending on its length, an interval [*t* − Δ*t, t*] along the fitness history of an allele may contain a number of (perhaps nested) valleys of varying depth (Figure A1). However, rather than attempt to record each valley as an individual event, we simply define *s* as the maximum valley depth traversed along an interval (see Appendix). A peculiar feature of this approach is that, for small values of *s*, we can choose an interval length Δ*t* long enough that *P*(*s*, Δ*t*) is a decreasing function of Δ*t* (i.e., since larger values of *s* are more likely to occur on longer time scales). However, below we will be concerned mainly with large values of *s* and time scales Δ*t* that are far from this sort of turnover region.

## 3. Results

To obtain data for *P*(*s*, Δ*t*) in a reasonable period of time, we limit our study to chains with at most 16 amino acids (802,075 conformations unrelated by symmetry [36]). The fitness landscapes of longer chains will clearly differ, however, we expect that, within reason, results for somewhat longer chains (e.g., 32 amino acids) will be similar, given proper adjustments to the mutation rate per gene *μ* (see below).

We simulate protein evolution for different chain lengths (*L* ≤ 16), native folds (Figures S2 and S5), population sizes (*N* ≤ 10^3^), and mutation rates (*μN* ≤ 2). In each situation, we conduct replicate simulations in parallel on multiple processors of a high performance computer [37]. Each processor begins with a monomorphic population constructed from *N* copies of a gene encoding a randomly selected amino acid sequence. Each population is then equilibrated until an allele reaches fixation with *P_N_* > 0.75. After this point, alleles are sampled at random from a population every 64*N* generations and their histories are recorded. For the most computationally intensive problems (i.e., for the largest chain lengths and mutation rates), we are able to generate 10^5^ samples in about ten days using 128 processors. For chains with 12 amino acids (15,037 conformations), samples can be obtained much more rapidly, and results for chains of 12 and 16 amino acids are actually very similar (computer code and sample data are available from the authors on request).

Since effective population size varies substantially across mammalian species, it is important to ask whether simulations conducted for a particular population size can be used to estimate *P*(*s*, Δ*t*) for larger (or smaller) populations evolving at the same overall rate *μN*, as expected from diffusion theory [40]. To answer this question, we compared the scaled distributions *P*(*Ns* > *x*, Δ*t*) for population sizes *N* = 100, 200, 500 and 1000. For a fixed mutation rate *μN*, we find that plots of *P*(*Ns* > *x*, Δ*t*) roughly collapse to the same curve. As a result, we can estimate *P*(*s*, Δ*t*) for realistic populations using a much smaller population size, which greatly reduces the amount of time spent on the simulations. The reason for this is fairly simple – the local structure of the fitness landscape, as measured by e.g. the distribution of fitness effects or the probability of compensatory neutral mutations, scales in a similar way with population size; As population size increases, the landscape in the neighborhood of an evolved sequence becomes less rugged in proportion to *N*.

Results of this exploration are described in Figures 1–2. In Figure 1, we plot the distributions of beneficial and deleterious fitness effects, *P*(Δ*w* > *x*) and *P*(−Δ*w* > *x*), respectively. Each pair of plots corresponds to a simulation for one of the population sizes listed above (the width of a plot increases with decreasing *N*). The results describe proteins with 12 amino acids folding to the native conformation in Figure S2, and the overall mutation rate in each simulation is *μN* = 1.

**Figure 1.**
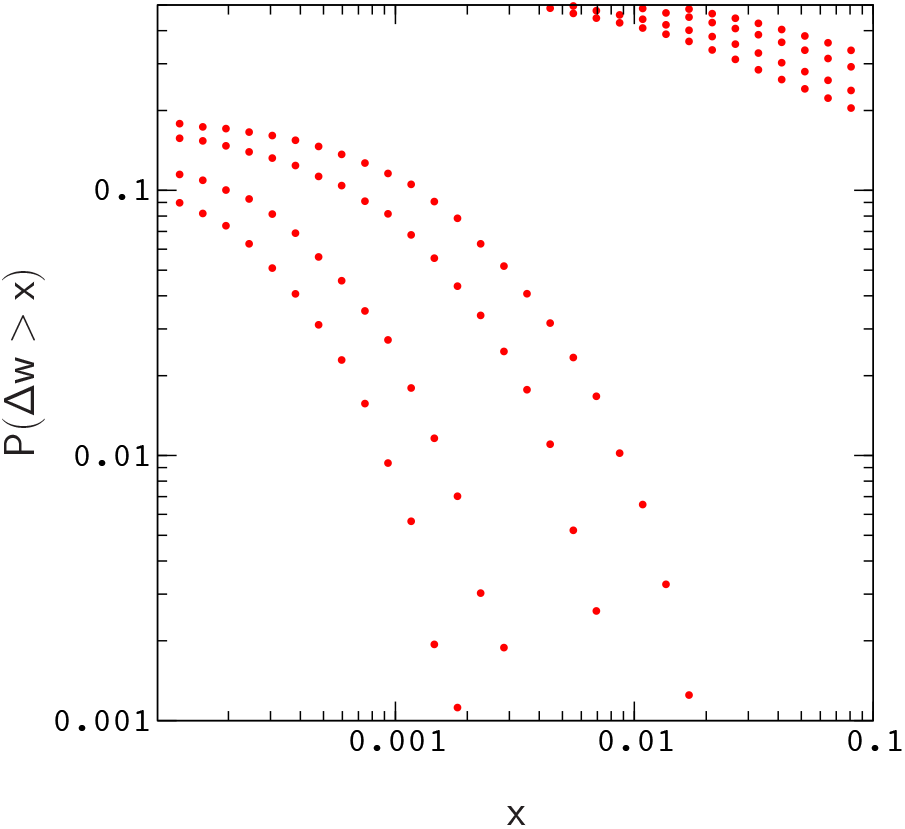
Distribution of beneficial fitness effects, *P*(Δ*w* > *x*). The distribution of deleterious fitness effects, *P*(−Δ*w* > *x*), is partially included in the upper right corner of the figure for reference. The plots are generated by randomly mutating evolved sequences sampled from simulations with *N* = 100, 200, 500, and 1000 (the width of a plot increases with decreasing *N*). If the data for Δ*w* in each plot is rescaled by the appropriate factor of *N*, the distributions *P*(*N*Δ*w* > *x*) roughly collapse to a single curve. For *N* = 1000, about 47 percent of the mutations are strongly deleterious (*N*Δ*w* < −5), about 28 percent are nearly neutral (−1 < *N*Δ*w* < 1), and about 0.7 percent are beneficial (*N*Δ*w* > 1), consistent with results obtained by Tamuri et al. [38] for mammalian mitochondrial proteins (Tamuri et al. use logarithmic fitness differences in their work, however, this distinction can be neglected when ln(1 + *x*) ≃ *x* [39]).

To generate data for Figure 1, we sampled the landscape around evolved genes using a simple procedure that mimics error–prone polymerase chain reactions [41]; The procedure begins from a large number of copies of an evolved gene. A Poisson random number of random mutations are applied to each copy, and the results are sorted by the number of mis–sense mutations (i.e., neglecting back mutations). For each simulation, we randomly selected 10^2^ evolved sequences. Each evolved sequence was then used to generate 10^4^ random single amino acid mutants, for a total of 10^6^ mutants per plot.

As is evident by closer inspection of Figure 1, the probability of a beneficial (or deleterious) mutation with effect Δ*w* > *x* decreases almost linearly with increasing population size; The scaled distributions *P*(*N*Δ*w* > *x*) for different population sizes roughly collapse onto a single curve. A similar result is obtained for the distribution of compensatory neutral double mutants *P*(*s* > *x*) (Figure S3). The collapse is shown explicitly for the valley crossing probability *P*(*s* > *x*, Δ*t*) in Figure 2.

**Figure 2.**
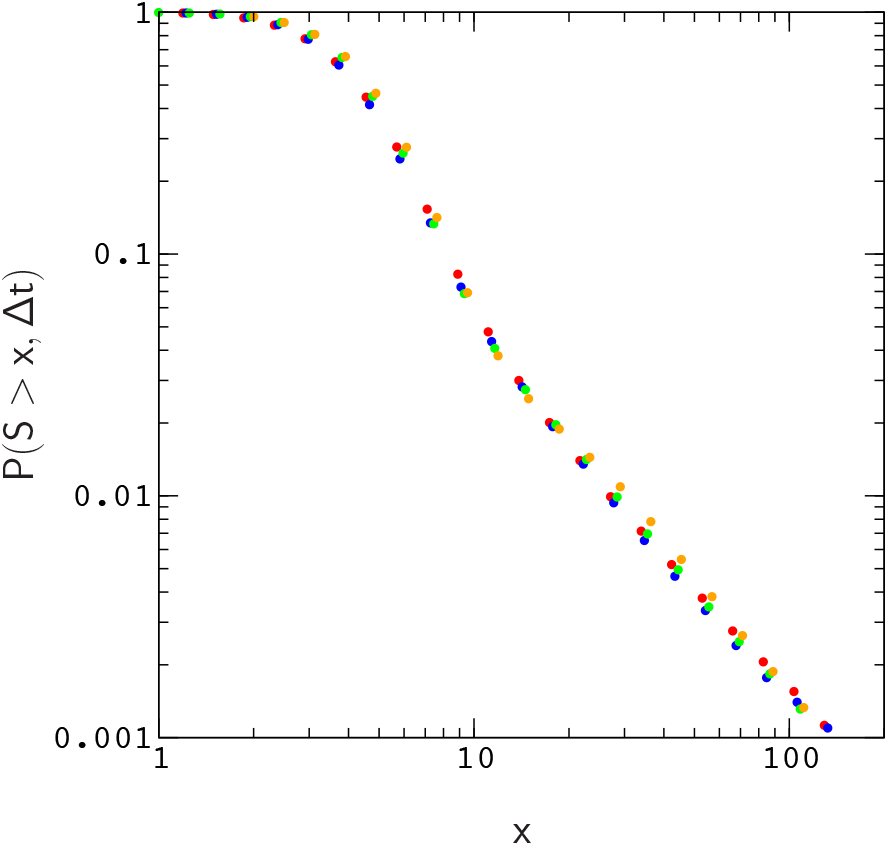
Probability, *P*(*S* > *x*, Δ*t*), that an allele, randomly sampled at time *t*, has crossed a valley of depth *S* = 2*Ns* in the interval [*t* − Δ*t, t*] for Δ*t* = 128*N* and *μN* = 1. The data describe the same model as in Figure 1. Plots for population sizes *N* = 100, 200, 500, and 1000 are colored red, blue, green, and orange, respectively.

Given this result, we now restrict our attention to populations with *N* = 200 and proteins with 16 amino acids. To compare our results to those of Meer et al., we require the site mutation rate in our model to agree with the pedigree rate for the control region in human mitochondria used in their estimate for mammalian mt tRNA molecules – about 1 × 10^−6^ per site per year. Assuming a typical length of about 80 nucleotides for tRNA molecules, a generation time of 20 years, and a (mitochondrial) effective population size of *N* = 2,500, we arrive at an overall mutation rate of *μN* ≃ 4 for human mt tRNA genes. To obtain the same site mutation rate for protein genes with 48 nucleotides (16 amino acids), we need an overall mutation rate of about *μN* ≃ 2.

We plot *P*(*S* > *x*, Δ*t*) versus Δ*t* for this situation in Figure 3, where *S* = 2*Ns*. The range of the plot, Δ*t* ≤ 128*N*, roughly corresponds to the time scale of human evolution (about six million years). Over this time scale, *P*(*S* > *x*, Δ) is roughly linear in Δ*t* for *x* ≳ 10. The time scale for mammalian evolution is much longer (on the order of tens of millions of years), however, on human time scales, the frequencies of large events in our model are already approaching the ten percent levels observed for Watson–Crick switching events in mammalian mt tRNA phylogenies [13]. For example, the probability of sampling an allele that has crossed a fitness valley of depth *S* > 9.1 for Δ*t* = 128*N* is about 0.36, the probability for *S* > 17.8 is about 0.07, and the probability for *S* > 27.8 is already about 0.03.

**Figure 3.**
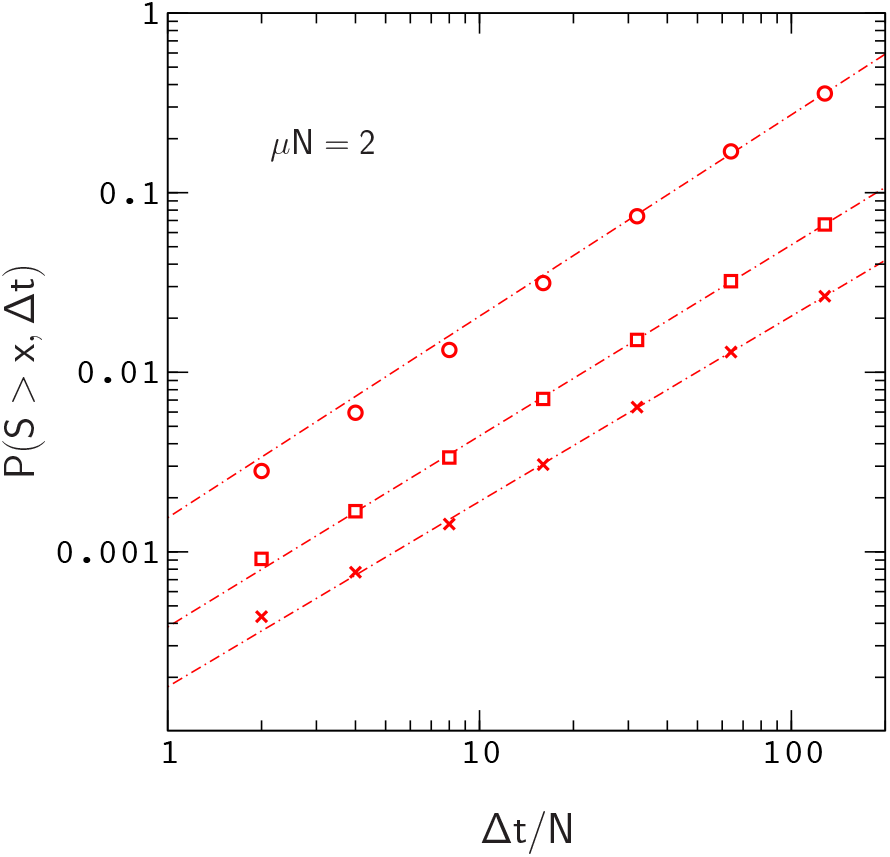
*P*(*S* > *x*, Δ*t*) versus Δ*t* for *x* ⋍ 9.1 (circles), 17.8 (squares) and 27.8 (crosses). The results describe proteins with 16 amino acids folding to the native conformation in Figure S5. Each data point is determined from over 10^5^ allele histories. The range of the plot, Δ*t* ≤ 128*N*, corresponds to the time scale of human evolution (about six million years). The dashed lines (power law fits to the data) are very close to linear, increasingly so for larger values of *x* (see Appendix for more details).

Clearly, these numbers will continue to increase for larger values of Δ*t* and larger mutation rates *μN* (Figure 4). In addition, *P*(*S* > *x*, Δ) will also increase with chain length since, for a constant site mutation rate, the mutation rate per gene *μ* is proportional to chain length. If we assume that the fitness landscapes of proteins with 16 amino acids can be used to represent the landscapes of larger chains, then e.g. doubling the mutation rate per gene will have the same effect as doubling the chain length to obtain a protein encoded by the same amount of genetic material as a tRNA molecule or small protein motif [42]. This is not an unreasonable assumption, since, as we have noted earlier (see the captions to Figures 1 and S3), the local structures of fitness landscapes in the model, as measured by the scaled distribution functions *P*(*N*Δ*w* > *x*) and *P*(*Ns* > *x*), are already similar to those inferred from real proteins with much longer sequences. In this case, extrapolating from the data in Figure 4, we find that the probability of sampling an allele that has crossed a valley of depth *S* > 27.8 over a time interval Δ*t* = 128*N* increases to about *P* ≃ 0.1. Thus, even neglecting the anticipated increase in *P*(*S* > *x*, Δ) for Δ*t* > 128*N*, the results for small protein motifs are already consistent with those of Meer et al.

**Figure 4.**
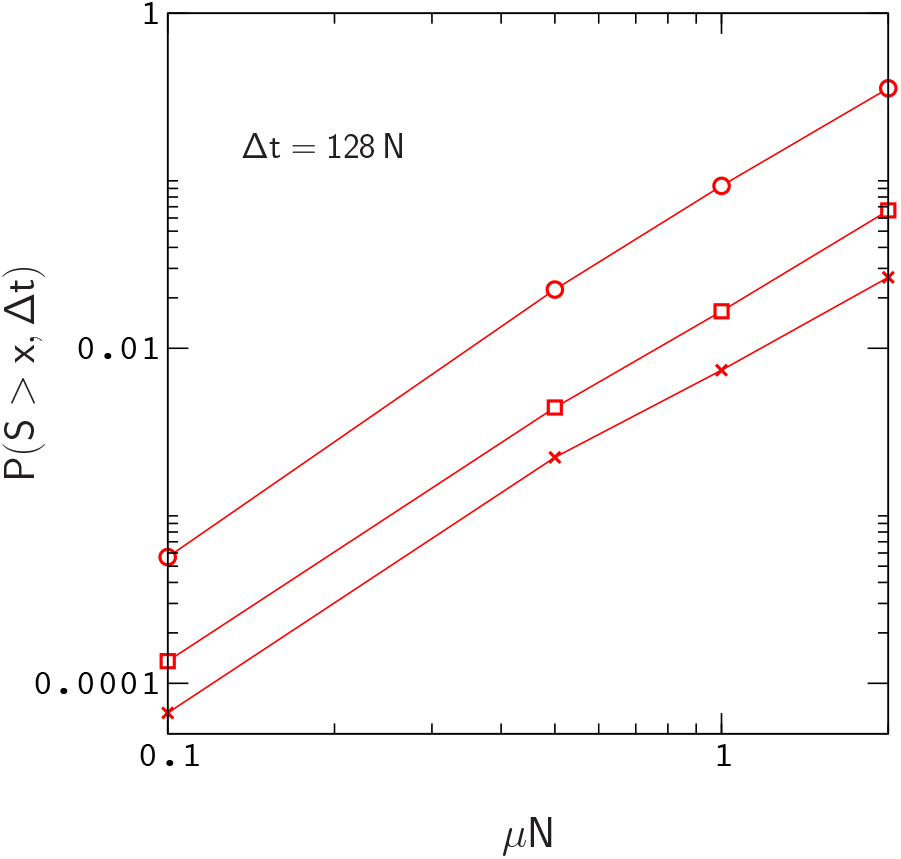
*P*(*S* > *x*, Δ*t*) versus *μN* for *x* ≃ 9.1 (circles), 17.8 (squares) and 27.8 (crosses). The results describe the same simulation data as Figure 3.

As a final note, it is important to remark that the deepest valleys navigated by alleles in our simulations actually correspond to events in which a deleterious mutation is, to a major extent, compensated by a mutation back to a similar amino acid type at the same site (see Figure S6). It is also worth noting that local fitness peaks sampled at various points in our simulations (by steepest ascent in fitness, starting from a randomly sampled sequence) are often separated by deep fitness valleys, or ravines.

## 4. Discussion

Does the model provide an accurate picture of fitness valley crossing for small protein motifs? This question is very difficult to answer due to the extreme complexity of protein fitness landscapes, and the unknown effects of varying host genetic backgrounds experienced by protein genes. However, from basic principles, it appears that the model is roughly accurate: Local features of protein fitness landscapes, such as the distribution of fitness effects, can be inferred from protein sequences by fitting a population dynamics model to branches of a phylogenetic tree [38] such that background effects are accounted for by allowing the parameters of the model to vary among branches. Tamuri et al. have used this type of approach to estimate the distribution of fitness effects for mammalian mitochondrial proteins [38], and our results for *P*(*N*Δ*w* > *x*) are in good agreement with their data (Figure 1). Given the similar nature of fitness conditions in each system (i.e., that proteins are also polymers required to fold into a specific shape in order to function), it is seems reasonable to expect that the topographies of model fitness landscapes resemble those of small protein motifs.

Longer chains with more developed core structures, and more restrictive fitness conditions such as binding to proteins within a larger domain, may lead to qualitatively different results due to the potential for larger compensatory effects [27], and the suppression of substitution rates in the core and binding interface regions. In addition, much larger mutation rates can occur in micro–organisms such as viruses [43], and our results suggest that, under these conditions, fitness valley crossing will be more pronounced. RNA viruses such as HIV–1 are also subject to high rates of recombination [44,45], which may interfere with valley crossing [5,6]. Because the computational cost of our simulations increases in proportion to the number of mutations (i.e., the number of fitness calculations), a full study of the model for virus proteins may be challenging. However, we hope to address these problems in future work.

## Supplementary Materials

The following are available online at http://www.mdpi.com/2073-4425/xx/1/5/s1, Figure S1: Plot of the logistic fitness function defined in Equation (3); Figure S2: Native fold studied in Figures 1–2; Figure S3: Plot of *P*(*s* > *x*) for the native fold studied in Figures 1–2; Figure S4: Plot of *P*(*S* > *x*) for an inferred Potts model of cytochrome *c* oxidase subunit 2; Figure S5: Native fold studied in Figures 3–4; Figure S6: Plot of *P*(*S* > *x*, Δ*t*) for the subset of deleterious mutations compensated by a mutation back to a similar amino acid type at the same site.

## Author Contributions

Conceptualization, Erik Nelson; Funding acquisition, Nick Grishin; Investigation, Erik Nelson; Software, Erik Nelson; Supervision, Nick Grishin.

## Acknowledgments

It is a pleasure to thank Daniel Moser for help with the computations, and two anonymous reviewers for helpful suggestions.

## Appendix

**Figure A1.**
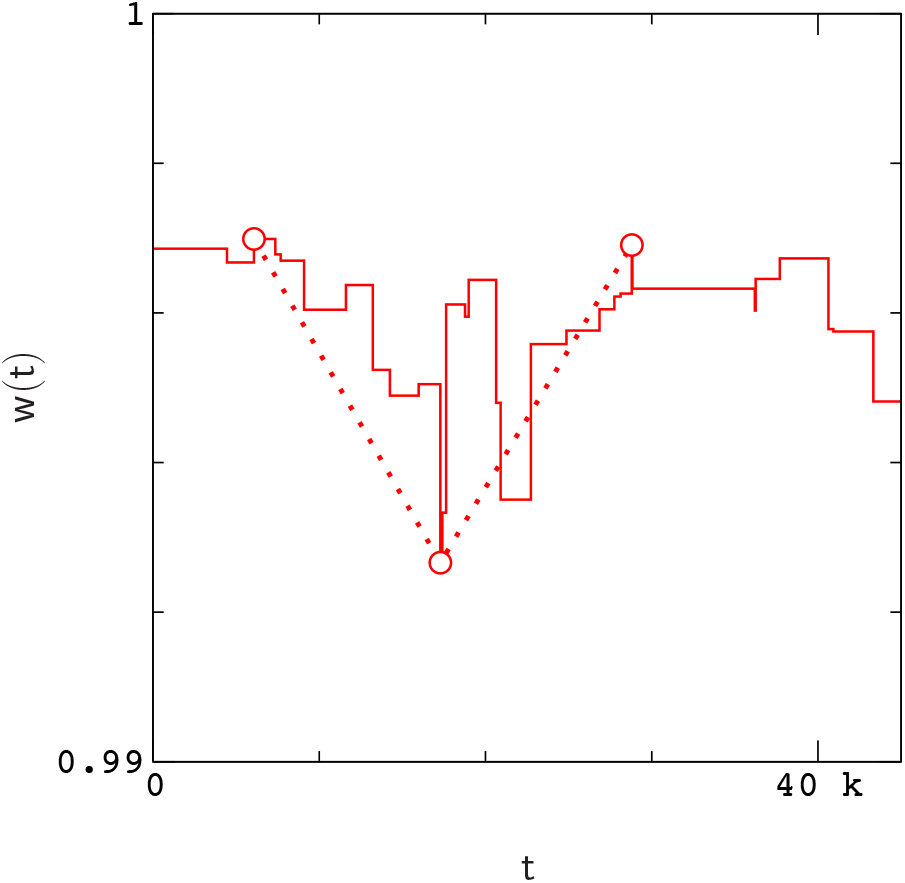
Fitness history of an allele between adjacent fixation events (solid line) sampled from a simulation with *N* = 10^3^ and *μN* = 1. Circles and dashed lines indicate the configuration of points which maximize the valley depth *s* = **min**{*w_i_* − *w_j_*, *w_k_* − *w_j_*}.

To compute the maximum valley depth traversed along an interval (Figure A1), we consider all possible placements of three points, *w_i_*, *w_j_* and *w_k_*, where *i* < *j* < *k* denote the occurrence times of mutations. For a given placement of points, valley depth s is defined as the smaller of the two fitness differences *w_i_* − *w_j_* or *w_k_* − *w_j_*. The maximum valley depth can then be expressed as,

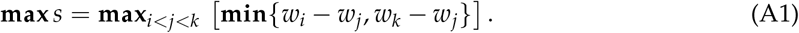

To avoid complicating our expressions, we use the plain symbols *s* and *S* = 2*Ns* to denote maximum valley depth in *P*(*s* > *x*, Δ*t*) and *P*(*S* > *x*, Δ*t*).

The structure of *P*(*S* > *x*, Δ*t*) can be explained roughly as follows: As we noted earlier, an interval [*t* − Δ*t*, *t*] along a fitness trace may contain a number of distinct valleys of varying depth. Valleys of large depth are rare on the time scale of human evolution and short lived. As a result, when *x* is large, doubling Δ*t* doubles the probability that a valley with depth greater than *x* will occur within an interval [*t* − Δ*t*, *t*], and *P*(*S* > *x*, Δ*t*) increases linearly with Δ*t*. This will be true as long as Δ*t* is not too large or too small; For large enough Δ*t*, *P*(*S* > *x*, Δ*t*) will begin to saturate (i.e., *P* → 1), at which point the slope of the curve, *∂P*(*S* > *x*, Δ*t*)/*∂*Δ*t*, tends to zero, and linearity is lost. Conversely, for small enough Δ*t*, the typical duration of an event (valley of depth greater than *x*) will begin to exceed Δ*t*, and again *∂P*(*S* > *x*, Δ*t*)/*∂*Δ*t* will begin to change. This change will depend both on *x* and the topography of valleys with depth greater than *x*, however, we have not explored this issue in detail.

