## Supplementary Material for "How often do Protein Genes Navigate Valleys of Low Fitness?"

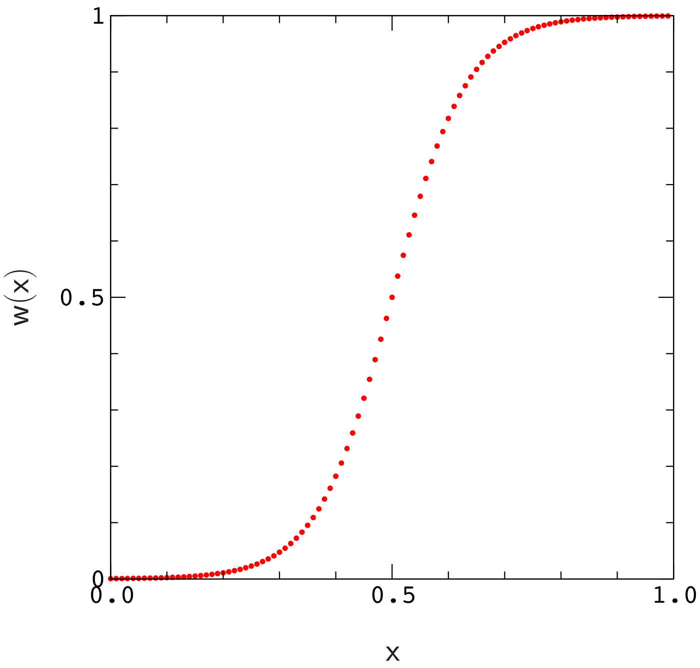


Figure S1. Plot of the logistic fitness function defined in Equation (3) for $k = 15$.

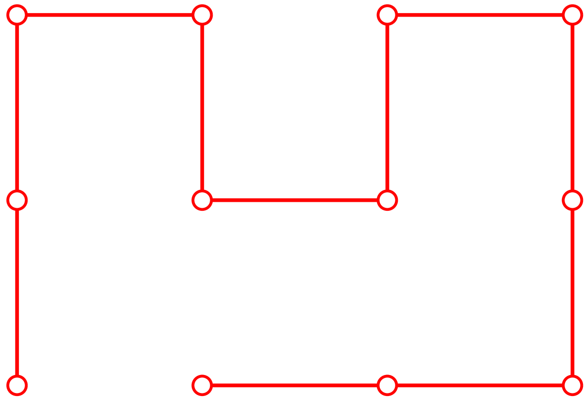

Figure S2. Native conformation studied in Figures 1–2.

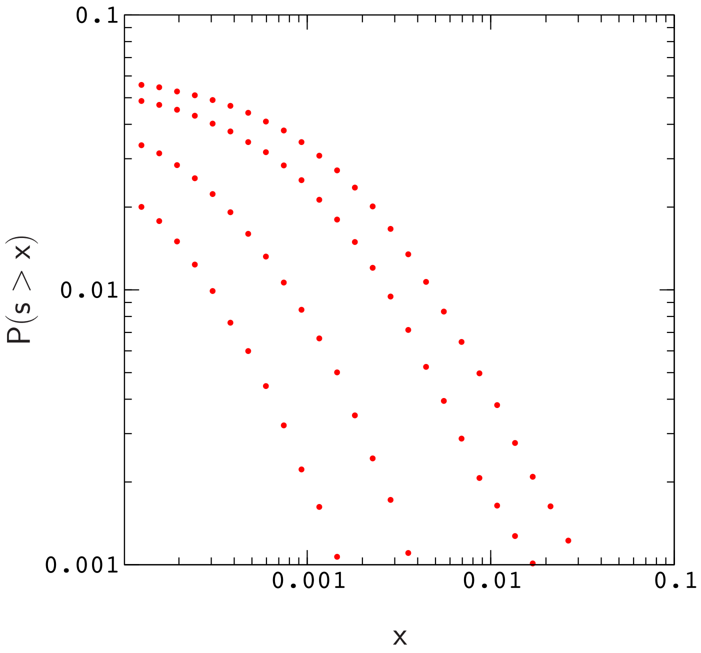


Figure S3. Frequency $P(s > x)$ of neutral, or beneficial double mutants with valley depth $s$. A double mutant is considered neutral or beneficial when its net effect $\Delta w$ satisfies $\Delta w > -1/N$; Valley depth is measured as described in the Appendix. The model, population sizes, and the procedure used to generate double mutant data for the plots are the same as in Figure 1 (the width of a plot increases with decreasing $N$). If the data for $s$ in each plot is rescaled by the appropriate factor of $N$, the distributions, $P(Ns > x)$, roughly collapse to a single curve. The results for $P(Ns > x)$ are consistent with results obtained using a Potts fitness model inferred from mitochondrial proteins in our most recent work (see Figure S4).

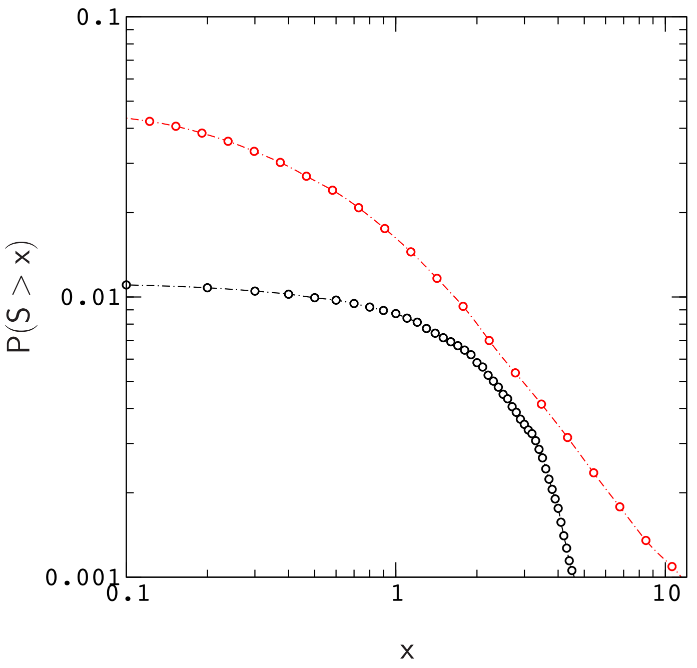


Figure S4. Frequency $P(S > x)$ of neutral or beneficial double mutants with valley depth $S = 2Ns$ for proteins folding to the conformation in Figure S5 (red), and for a Potts fitness model of fish cytochrome c oxidase subunit 2 (black). A double mutant is considered neutral or beneficial when its net effect $\Delta w$ satisfies $\Delta w > -1/N$, and valley depth is measured as described in the Appendix. The procedure used to generate double mutant data for these plots is the same as that used in Figure 1 (for cytochrome c oxidase 2, amino acid mutations are restricted to the "predictive subset", $p_i(a) \geq 1/\sqrt{B}$, where $p_i(a)$ is the probability of finding an amino acid type $a$ at site $i$ in the alignment used to construct the Potts model, and $B$ is the number of sequences in the alignment. This form of restriction is also applied to double mutants generated for the model).

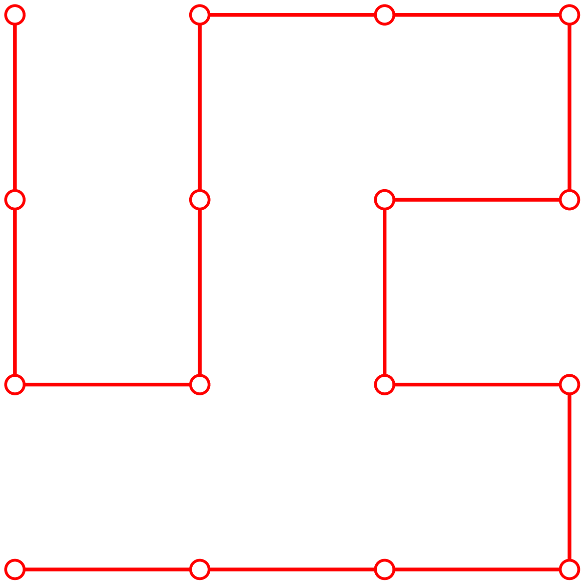

Figure S5. Native conformation studied in Figures 3–4.

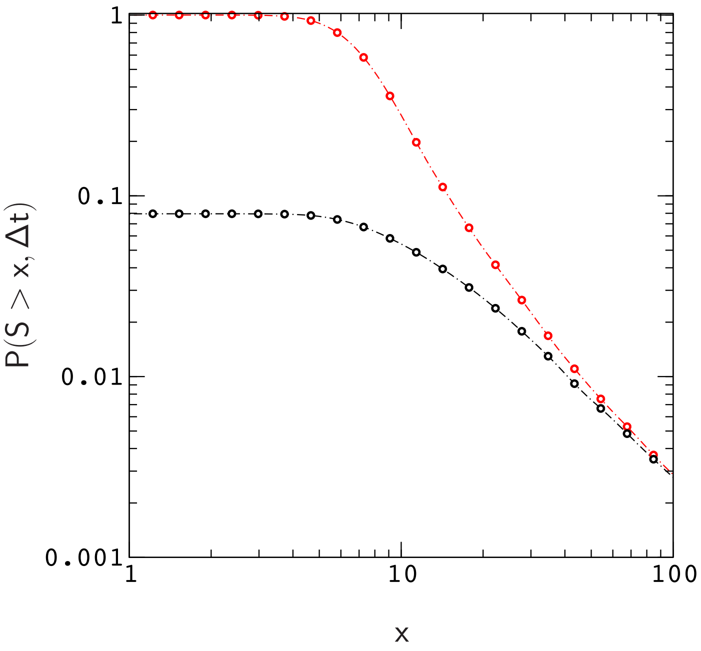


Figure S6. $P(S > x, \Delta t)$ versus $x$ for all allele histories (red) and histories in which more than ninety percent of the valley depth obtained in Equation A1 is due to a deleterious mutation followed by a compensatory mutation at the same amino acid site (black). The plots describe a simulation of proteins folding to the conformation in Figure S5 with $N = 200$, $\mu N = 2$, and $\Delta t = 128N$.
